# The role of alpha-helix on the structure-targeting drug design of amyloidogenic proteins

**DOI:** 10.1101/2020.11.20.391409

**Authors:** Carmelo Tempra, Carmelo La Rosa, Fabio Lolicato

## Abstract

The most accredited hypothesis links the toxicity of amyloid proteins to their harmful effects on membrane integrity through the formation of prefibrillar-transient oligomers able to disrupt cell membranes. However, damage mechanisms necessarily assume a first step in which the amyloidogenic protein transfers from the aqueous phase to the membrane hydrophobic core. This determinant step is still poorly understood. However, according to our lipid-chaperon hypothesis, free lipids in solution play a crucial role in facilitating this footfall. Free phospholipid concentration in the aqueous phase acts as a switch between ion channel-like pore and fibril formation, so that high free lipid concentration in solution promotes pore and repress fibril formation. Conversely, low free lipids in the solution favor fibril and repress pore formation. This behavior is due to the formation of stable lipid-protein complexes. Here, we hypothesize that the helix propensity is a fundamental requirement to fulfill the lipid-chaperon model. The alpha-helix region seems to be responsible for the binding with amphiphilic molecules fostering the proposed mechanism. Indeed, our results show the dependency of protein-lipid binding from the helical structure presence. When the helix content is substantially lower than the wild type, the contact probability decreases. Instead, if the helix is broadening, the contact probability increases. Our findings open a new perspective for in silico screening of secondary structure-targeting drugs of amyloidogenic proteins.

## Introduction

Membranes surrounding living cells are soft interfaces made up of thousands of different lipids species crowded with numerous membrane-associated proteins (Dupuy and Engelman, 2008; Sezgin et al., 2017). Membrane-protein and protein-protein interactions are responsible for a wide-ranging of cellular functions, including signaling, transport, energy storage and the catalysis of essential biochemical reactions (Krogh et al., 2001; Overington et al., 2006; Sezgin et al., 2017). Many proteins are mostly unstructured in solution and are described as intrinsically disordered (IDP) (Chiti and Dobson, 2017). Some of these proteins are amyloidogenic, forming beta-sheet rich fibrillar insoluble aggregates through a series of conformational transitions (amyloid cascade) (Selkoe and Hardy, 2016). Such aggregates have been linked with human diseases since large aggregates containing amyloidogenic proteins were found in the brain and pancreas (amyloid hypothesis). However, the correspondence between fibrillar plaques and diseases is not always satisfied. (Sengupta et al., 2016; Westermark, 2011). An increasing number of reports link the toxicity of amyloid proteins to their harmful effects on membrane integrity, but the molecular mechanism underlying amyloidogenic proteins transfer from the aqueous environment to the membrane’s hydrocarbon core is poorly understood. Recently, the most accredited hypothesis of the toxicity suggests the toxic species are the prefibrillar-transient oligomers (Benilova et al., 2012; La Rosa et al., 2020; Lee et al., 2017; Schemmert et al., 2019) which form ion channel-like pores into the membrane (Chiti and Dobson, 2017; Quist et al., 2005; Scalisi et al., 2010; Sciacca et al., 2012; Scollo and La Rosa, 2020). Three major membrane damage models have been proposed so far: (i) generation of stable transmembrane protein pores (the “toxic oligomer hypothesis”); (ii) membrane destabilization via a “carpet model” or iii) removal of lipid components from the bilayer by growing amyloids (the “detergent-like mechanism”). However, both of these damage mechanisms necessarily assume a first step in which the amyloidogenic protein transfers from the aqueous phase to the bilayer’s hydrocarbon core; this determinant step is poorly understood. Whether these three models are mutually exclusive or if (and how) they cooperate in triggering membrane damage remains to be established.

For these reasons, understanding the early steps of the transfer of proteins from the water environment into the lipid wall is of vital importance. According to our lipid-assisted protein transport model (La Rosa et al., 2016), water-soluble lipid-protein complexes insert into the membrane faster than the bare protein, provided the hydrophobicity of the lipid-protein complex is higher than that of the bare protein. This model, called the lipid-chaperon hypothesis, was tested on human and rat amylin (IAPP), Aβ, α- and β-synuclein (Sciacca et al., 2020; Scollo et al., 2018). Free phospholipid concentration in the aqueous phase acts as a switch between ion channel-like pore and fibril formation, so that high free lipid concentration in solution promotes pore and repress fibril formation. Conversely, low free lipids in the solution favor fibril and repress pore formation. This behavior is due to the formation of stable lipid-protein complexes.

Moreover, circular dichroism (CD) spectroscopy experiments highlight the presence of an alpha-helix structure when proteins interact with free-lipids. Indeed, CD spectra of Aβ_1-40_ showed a shift characteristic of helical structures when protein is incubated with 1,2-dimyristoleoyl-sn-glycero-3-phosphocholine lipid (PC14) at its own critical micelle concentration (CMC). In absence of PC14 Aβ_1-40_ CD spectra showed, instead, the characteristic beta-sheet structure. This finding strongly points out the role of small alpha-helix structured oligomers to be responsible for the toxicity on the membrane which is in agreement with thermodynamic and kinetic models considering pore formation through helical structures (Engel et al., 2006; Pannuzzo et al., 2013; Sparr et al., 2004). All in all, this agree with the general model where the random coil and β-sheet propensity are favorable in water (Apetri et al., 2006; Shea et al., 2019; Uversky, 2019) and α-helix are favorable in a membrane-like environment (Pannuzzo et al., 2013; Sparr et al., 2004).

Along with the CD experiments, we employed molecular dynamics (MD) simulations to get atomistic details on the protein/lipid interaction, identifying which amino acids showed more affinity for the lipid (Sciacca et al., 2020). Here, we hypothesize that the helix propensity is a fundamental requirement to fulfill the lipid-chaperon model. The alpha-helix region binds strongly with amphiphilic molecules (free lipids, oxidized lipids, free fatty acids), fostering the proposed mechanism (**Figure1 A**). Starting from our previous study, we employed classical MD simulations to assess helix propensity’s influence in the binding between amyloidogenic proteins (Aβ_1-40_ and hIAPP) and lipids (DEPC). Here, we demonstrate that the helix structure’s destabilization, achieved by introducing point mutations, impairs the binding with free lipids. Our study confirms the critical role of the alpha-helix structure in the binding with amphiphilic molecules suggesting new strategies to inhibit the cytotoxicity of amyloidogenic proteins (**Figure 1 B-C**).

**Figure 1.**
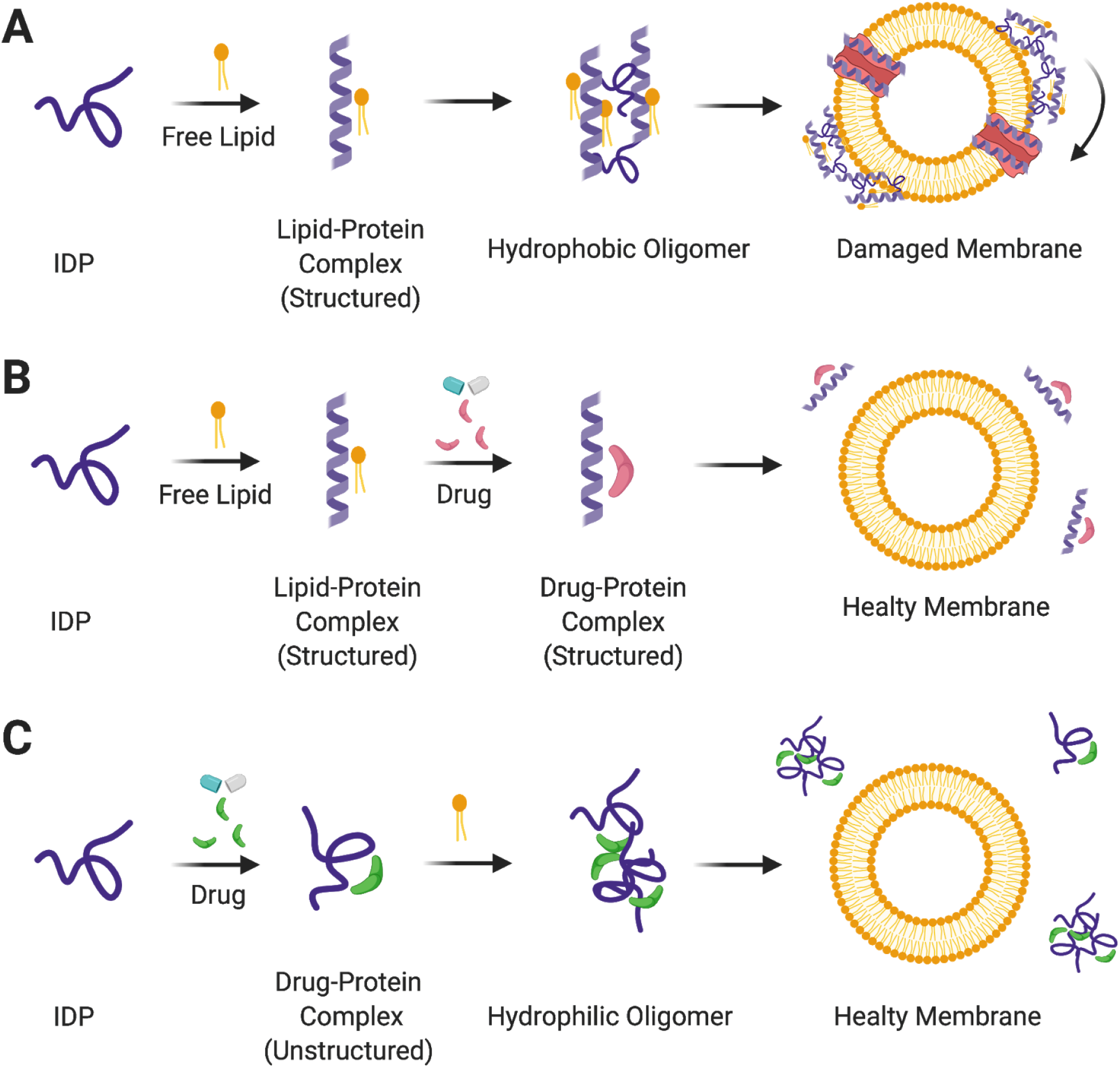
The alpha-helix region binds strongly with amphiphilic molecules (free lipids, oxidized lipids, free fatty acids), fostering the formation of hydrophobic structured oligomers able to damage the membrane (panel A). Our model suggests new strategies to inhibit the cytotoxicity of amyloidogenic proteins by designing structured-targeting drugs that can impair oligomer formation by competing with free lipids in binding with the alpha-helix structure (B) or preventing the helix structure formation (C). Created with BioRender.com.

## Results and Discussion

To shed light on the role of helical propensity in stabilizing the binding with free lipids in solution, we studied, computationally, how the substitution of critical amino acids, by point mutations, affects the secondary structure of hIAPP and Aβ_1-40_ and their binding with DEPC free lipid. The selected mutations were chosen by analyzing the contact occupancy between DEPC and the two amyloidogenic proteins, as reported in our previous study (Sciacca et al., 2020). The four amino acids with the highest contact occupancy were selected from the helical regions and mutated to Proline. As shown in **Figure 2**, we mutated H13, H14, L17, F20, and Y9, L12, F15, L16 for Aβ_1-40_ (left panel) and hIAPP (right panel), respectively. We also built a system containing all the single mutations at once. Proline is known to be a secondary structure breaker (Chou and Fasman, 2006; Kabsch and Sander, 1983; Li et al., 1996) and the substitution with Proline into Aβ_1-40_ and hIAPP aims to destabilize the helix propensity. Our goal is to link the loosing of the helical structure to the strength of the binding with the lipid.

**Figure 2.**
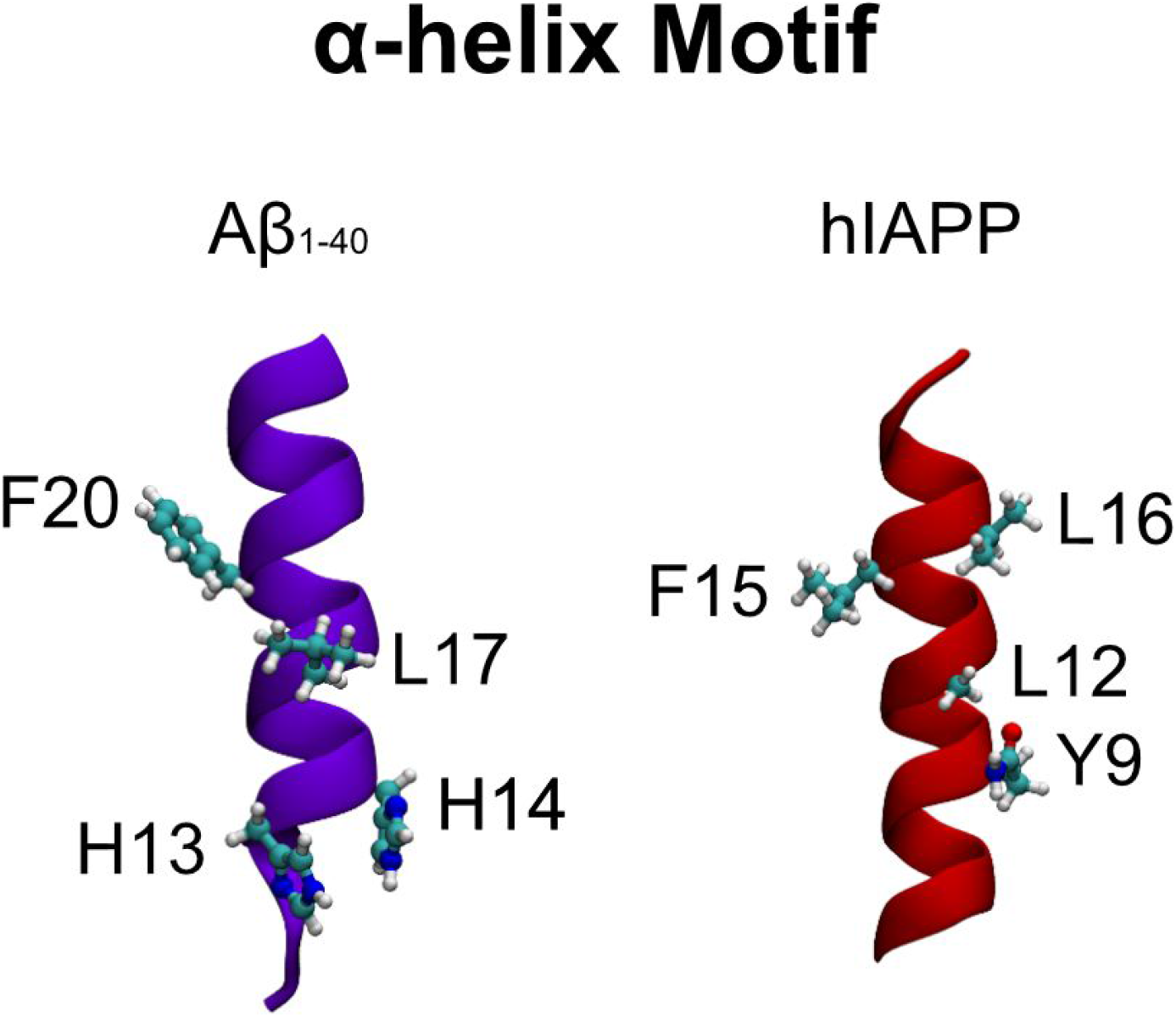
α-helix motif of Aβ_1-40_ (left panel) and hIAPP (right) proteins.

All the mutated proteins were first stabilized in water for one microsecond, in triplicate replica, using the CHARMM36m forcefield (Huang et al., 2017). The final structure of each simulation was extracted and randomly placed in a box containing a DEPC molecule. More details about the simulations can be found in the Methods section and Table 1.

We calculated the average amino acid-DEPC occupancy of contact using gmx mindist tool (Abraham et al., 2015) and in-house python script to compare the strength of the binding between each mutated protein and free lipid. **Figure 3 A** shows the contact occupancy for Aβ_1-40_ (left panel) and hIAPP (right panel), respectively. Both Aβ_1-40_ and hIAPP wild type proteins show a clear interaction motif at residues 9-16 and 13-20, respectively (top row). For Aβ_1-40_, the mutation at position 20 (Aβ_1-40_-F20P) drastically affects the binding with lipid, weakening the interaction with the 13-20 motif. All the other single mutations seem to not interfere much with the binding with the lipid. hIAPP case is, however, different. Only the hIAPP-F15P mutant shows a drastic decrease in the protein’s helix continent (**Figure 3B**). Interestingly, in all cases, the single mutant forms show an increase in the number of contacts. The interaction pattern is no longer limited to the 9-16 motif, but it is spread to a more broad amino-acid range.

**Figure 3.**
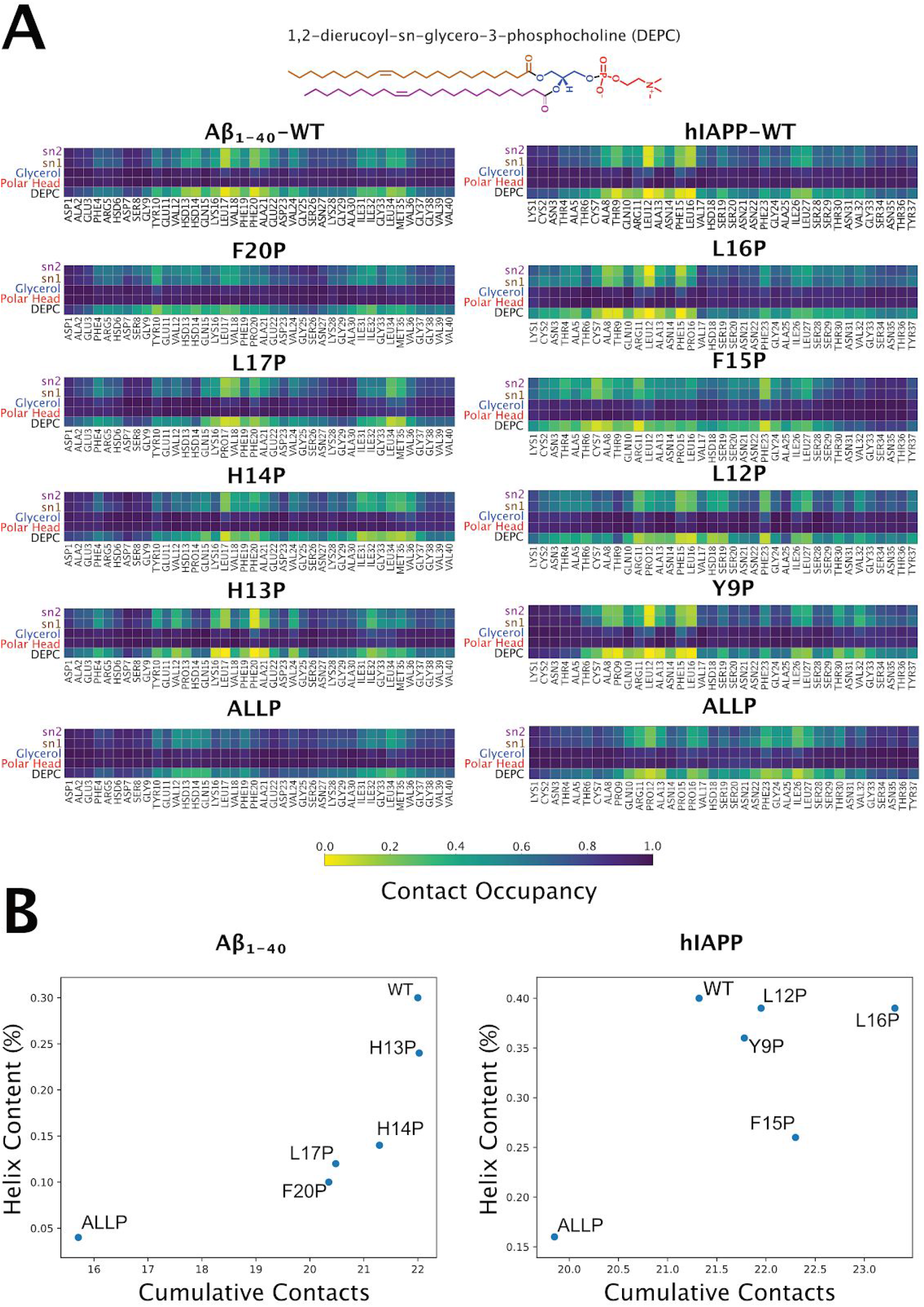
**Panel A** shows the pairwise contact occupancy map for Aβ1-40 - (left panel) and hIAPP- (right) DEPC complexes. Contact occupancy equal to 1.0 corresponds to the situation where a given interaction has taken place for the entire duration of the timeframe examined. The average was calculated by concatenating the last 500 ns from all three repeats. A contact is defined in any of the atoms between the two groups that were closer than 0.6 nm. **Panel B** shows, instead, the Helix content percentage in the function of the cumulative contacts for each of Aβ1-40- (left panel) and hIAPP- (right) DEPC complexes.

The simultaneous mutation of all four selected amino acids to Proline (Aβ_1-40_-ALLP and hIAPP-ALLP systems), induce, in both proteins, a drastic decrease in the number of contacts with the lipid (**Figure 3A**, last row) and the helix content (**Figure 3B** and **Figure 4**). **Figure 3B** shows, indeed, the behavior of the helicity content in the function of the total cumulative number of contacts. In the case of Aβ_1-40_, we observed a consistent trend: if the helicity is reduced, the total number of contacts with the lipids is drastically affected. As discussed above, the same trend is not, however, observed for the hIAPP, except for hIAPP-ALLP (where both helicity and contacts are drastically reduced). When a single point mutation reduces the helicity, the number of contacts increases. This different behavior between hIAPP and Aβ_1-40_ proteins can be explained by looking carefully into each mutation’s effect on the helix’s content (helicity) along the sequence. An in-depth look into the helix propensity per amino acid (**Figure 5**) highlights how mutations L12, F15, and L16 make the helix extension broader compared to the wild type. Even with a slight decrease in the total helix propensity, a broader helix extension helps gain contact affinity between protein and lipid. Indeed, contact maps (**Figure 3A-B**) show a broader contact range when helix propensity gets broader. In our precedent study (Sciacca et al., 2020), we were led from experimental observation to use helical structure to model protein/lipid interaction. Here we underline that helical structure is essential for the stability of the lipid/protein complex. In both proteins cases, the conclusion is the same: The lipid-protein interaction occurs only between the alpha-helix and the lipids, tails. This is following our model and recent studies, which indicates the lipid-protein complex’s hydrophobicity to be the culprit of membrane damage or fibrils formation.

**Figure 4.**
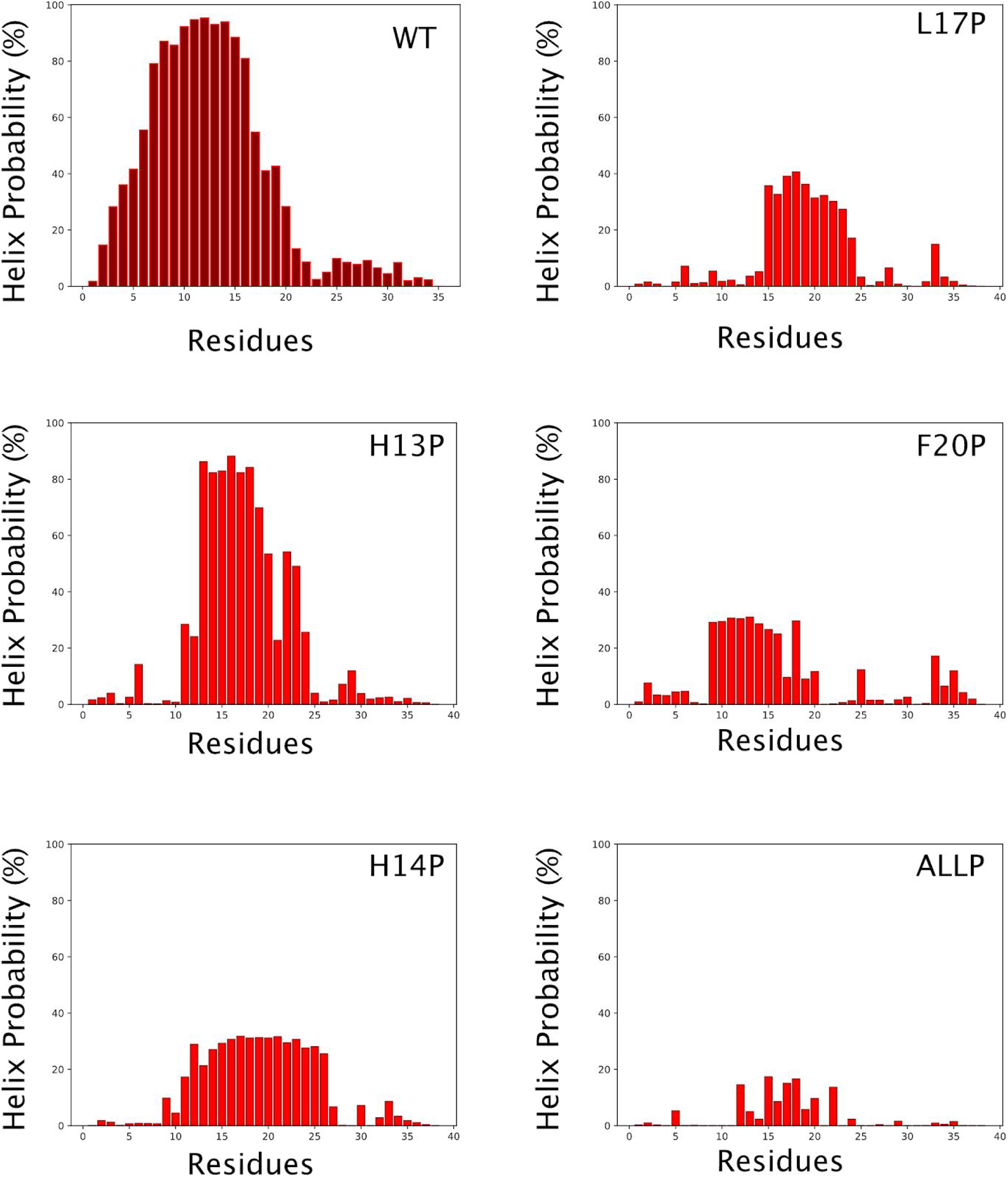
Residues-wise helix probability distribution for Aβ_1-40_ systems.

**Figure 5.**
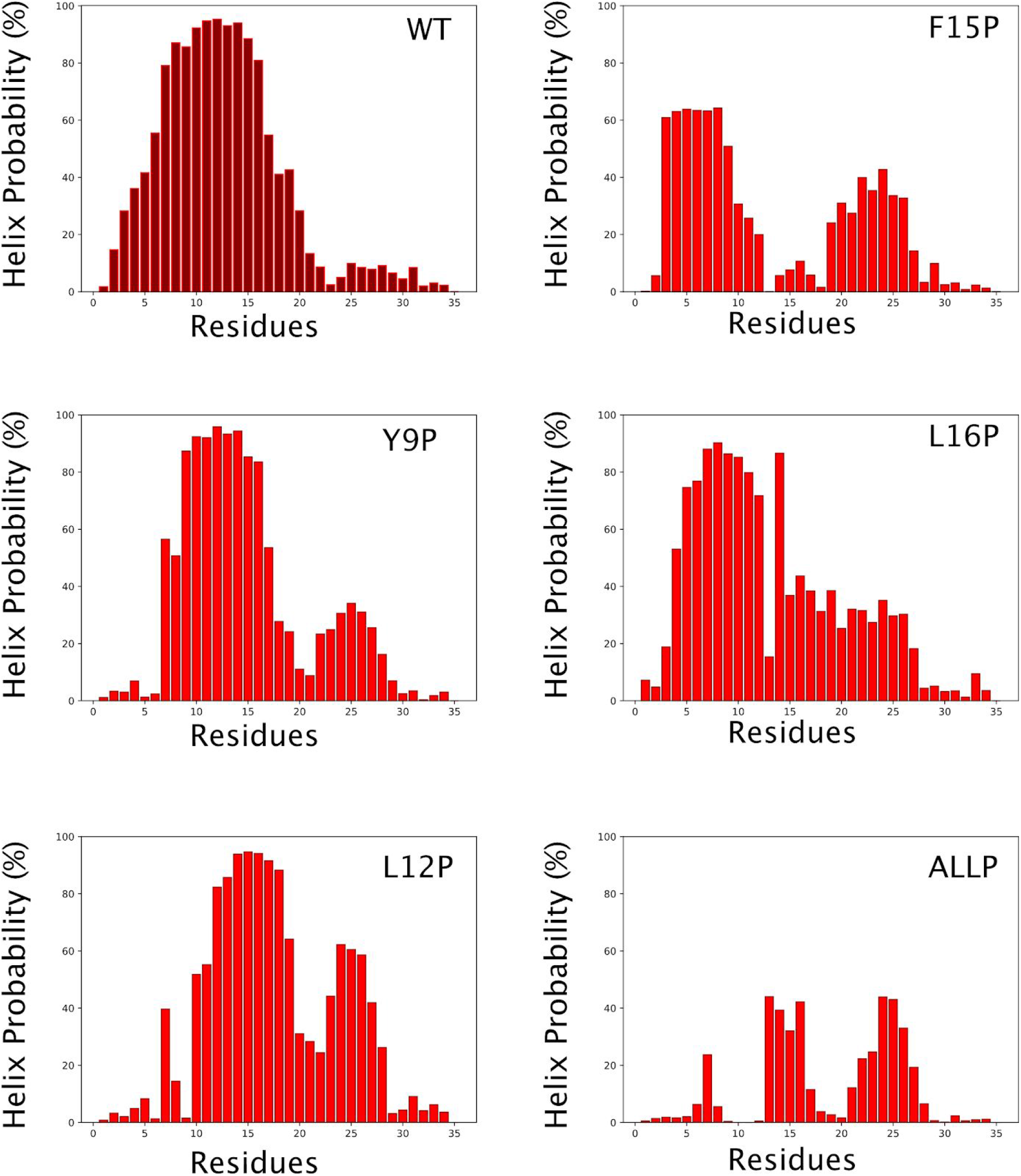
Residues-wise helix probability distribution for hIAPP systems.

## Conclusion

We introduce punctual proline mutations into Aβ_1-40_ and hIAPP to destabilize the helix propensity and check whether this can affect the contact affinity with the DEPC. The introduced mutations in Aβ_1-40_ were able to destabilize the helical structure, and a clear correlation between contact probability and total helix content was found, pointing out how the helix propensity is required for the protein/lipid complex stability. hIAPP, however, showed a certain toughness in losing helical structure, requiring introducing four mutations at one to decrease the helix content drastically. Our results show the dependency of protein-lipid binding from the helical structure presence. When the helix content is substantially lower than the wild type, the contact probability decreases. Instead, if the helix is broadening, the contact probability increases. To our knowledge, the lipid-chaperone mechanism plays an important role in the cytotoxicity of the proteins studied here. So far, an exciting challenge that has not been entirely addressed is the difficulty of successfully using docking and virtual screening methods aiming to block the cytotoxicity of amyloid peptides. The main problem could be related to the difficulty of having a well-defined structure to apply docking protocols. Indeed, using a random coil structure can be downright misleading when using docking protocols or molecular dynamics simulations. Here, we identified a structural motif that is crucial for explaining the chaperone-like mechanism, which is related to cytotoxicity. This helical motif can be used as a docking target to inhibit the lipid-chaperone mechanism and cytotoxicity in two ways. Firstly, drug molecules can be designed to compete with free lipids in solutions for the binding with amyloidogenic proteins (as illustrated in **Figure 1B**). Secondly, specific target molecules can be designed to destabilize the alpha helix region (**Figure 1C**). favoring the random coil structure and impairing the formation of hydrophobic aggregates responsible for the cytotoxicity.

## Methods

The selected mutations for hIAPP are Thr9, Leu12, Phe15, and Leu16, whereas Aβ_1-40_’s mutations are His13, His14, Leu17, and Phe20. Single point mutations plus a mutant containing all the mutations at once were done using CHARMM-GUI (Lee et al., 2016). The initial structure was taken from our previous simulations (Lolicato, F. and Tempra, C., 2020; Sciacca et al., 2020) both hIAPP (PDB ID: 2KB8 (Patil et al., 2009)) and Aβ_1-40_ (PDB ID: 1Z0Q (Tomaselli et al., 2006)) ‘s structures contain alpha helix. Each system was placed in cubic boxes containing water and counterions and simulated for one microsecond in a triplicate replica. Simulations details are listed in **Table 1**.

**Table 1.** Simulations details for protein systems.

| System | # water | # Na+ | # Cl- | # Replica | Time (μs) |
| --- | --- | --- | --- | --- | --- |
| A$\beta_{1-40}$-H13P | 11019 | 3 | 0 | 3 | 1 |
| A$\beta_{1-40}$-H14P | 11022 | 3 | 0 | 3 | 1 |
| A$\beta_{1-40}$-L17P | 11019 | 3 | 0 | 3 | 1 |
| A$\beta_{1-40}$-F20P | 11017 | 3 | 0 | 3 | 1 |
| A$\beta_{1-40}$-ALLP | 11026 | 3 | 0 | 3 | 1 |
| hIAPP-Y9P | 11037 | 0 | 2 | 3 | 1 |
| hIAPP-L12P | 11036 | 0 | 2 | 3 | 1 |
| hIAPP-F15P | 11042 | 0 | 2 | 3 | 1 |
| hIAPP-L16P | 11038 | 0 | 2 | 3 | 1 |
| hIAPP-ALLP | 11046 | 0 | 2 | 3 | 1 |

**Table 2.** Simulation details for protein/DEPC complex systems.

| System | # water | # Na+ | # Cl- | # Replica | Time (μs) |
| --- | --- | --- | --- | --- | --- |
| A$\beta_{1-40}$-H13P - DEPC | 6747 | 3 | 0 | 3 | 1 |
| A$\beta_{1-40}$-H14P / DEPC | 6747 | 3 | 0 | 3 | 1 |
| A$\beta_{1-40}$-L17P / DEPC | 6747 | 3 | 0 | 3 | 1 |
| A$\beta_{1-40}$-F20P / DEPC | 6747 | 3 | 0 | 3 | 1 |
| A$\beta_{1-40}$-ALLP / DEPC | 6747 | 3 | 0 | 3 | 1 |
| hIAPP-Y9P / DEPC | 6748 | 0 | 2 | 3 | 1 |
| hIAPP-L12P / DEPC | 6748 | 0 | 2 | 3 | 1 |
| hIAPP-F15P / DEPC | 6748 | 0 | 2 | 3 | 1 |
| hIAPP-L16P / DEPC | 6748 | 0 | 2 | 3 | 1 |
| hIAPP-ALLP / DEPC | 6748 | 0 | 2 | 3 | 1 |

The final structure of each simulation was extracted and randomly placed in a box containing a DEPC molecule. The box was solvated, neutralized, energy minimized, and simulated for one microsecond (see **Table 2** for simulation details). The following setup has been used for all the simulations. Leap-Frog integrator with a two femtosecond time step was used. Verlet algorithms with an updating frequency of 20 steps and a cut-off length of 1.2 nm have been employed. Particle Mesh Ewald algorithm with a cut-off length of 1.2 nm was used to calculate electrostatic interaction. Van der Waals interactions were treated employing a force-switching algorithm with a cut-off of 1.2 nm. To keep the temperature and pressure constant, Nose-Hoover thermostat and Parrinello-Rahman barostat coupling were used, respectively. The temperature was kept at 298 K with a coupling time of 1.0 ps, whereas the pressure was maintained at 1 bar with a coupling time of 5.0 ps. All covalent bonds with hydrogen were constrained using the LINCS algorithm.

## Acknowledgments

F.L. wishes to thank the CSC–IT Center for Science (Espoo, Finland) for computational resources.

## Data Availability

Simulations data are publicly available on Zenodo (Tempra, C., Lolicato, F., 2020) at the following address: 10.5281/zenodo.4280539

